# Two distinct cytokine response clusters identified in healthcare workers with apparent resistance to infection with *Mycobacterium tuberculosis* despite sustained occupational exposure

**DOI:** 10.1101/2025.03.27.643001

**Authors:** Avuyonke Balfour, Nomawethu Masina, Artur T. L. Queiroz, Bruno B. Andrade, Eduardo R. Fukutani, Abulele Bekiswa, David Lewinsohn, Deborah Lewinsohn, Rene Goliath, Katalin A. Wilkinson, Charlotte Schutz, Graeme Meintjes, Muki S. Shey

**Author notes:** Contributed equally to this manuscript (Avuyonke Balfour, Nomawethu Masina). **Corresponding Author** Muki Shey, Department of Medicine & CIDRI Africa, Faculty of health Sciences, University of Cape Town., Anzio Road, Observatory, Cape Town, South Africa, 7925.

## Abstract

**Introduction:** Evidence exists that some individuals resist (“Resisters”) *Mycobacterium tuberculosis* (Mtb) infection despite sustained exposure, but the protective mechanisms involved are not fully understood. We investigated immune responses induced by *ex vivo* stimulation of peripheral blood mononuclear cells (PBMC) with live Mtb in Resisters compared to those with latent TB infection (LTBI).

**Methods:** HIV-uninfected healthcare workers working in high TB exposure healthcare facilities for over 5 years were screened for Mtb sensitization using the interferon-gamma release assay (IGRA) and the tuberculin skin test (TST). We identified Resisters (TST<10mm and IGRA<0.35 IU/mL, n=129) and those with LTBI (TST≥10mm and IGRA≥0.35 IU/mL n=145). We selected a subset of ‘extreme resisters’ (TST=0mm and IGRA<0.2 IU/mL; n=26) and ‘extreme LTBI’ (TST≥15mm and IGRA≥1 IU/mL; n=24) for these analyses. Blood was collected and PBMC isolated and cultured with live H37Rv Mtb for 18 hours. Supernatants were collected and used for measuring 65 secreted analytes by Luminex. We also evaluated cell-associated cytokine expression by CD4 T cells and monocytes in these participants using flow cytometry.

**Results:** Using the feature selection in R, we identified a set of 4 cytokines: IL-17A, MCP-1, IL-8, and MDC, which collectively classified extreme Resisters from extreme LTBI with an area-under-the-curve (AUC) of 0.67 (0.54-0.85). Focusing only on the extreme Resisters, and using hierarchical clustering, participants in this group segregated into two main clusters (Resister_c1 and Resister_c2). Further analyses identified 37 cytokines that were significantly higher and 15 cytokines that were lower in Resister_c2 compared with Resister_c1. A set of 5 cytokines (TRAIL, MIP-1β, Fractalkine, GRO-α and IL-1α) collectively classified these two clusters with an AUC of 1 (1-1). CD4 T cell and monocyte responses to Mtb did not significantly differ between the two clusters.

**Conclusions:** Mtb-specific immune responses segregate extreme Resisters into two distinct clusters in individuals with apparent resistance to Mtb infection and may allow discrimination of individuals with true resistance from those with infection but have alternative immune responses not detected by IGRA and TST.

## Introduction

Sustained exposure of individuals with infectious TB results in latent TB infection (LTBI) in most people. Healthcare workers (HCW) have an increased risk of infection with Mtb and developing TB due to occupational exposure to TB patients. Despite sustained exposure, some HCW remain negative on tests for TB infection, suggesting they may be protected. Several previous studies have suggested that resistance to Mtb infection exists including household contact (HHC) studies from Uganda, multisite HHC study of MDR-TB patients, gold miners in South Africa, and recently our study on HCW in Cape Town [1-4]. While mechanisms associated with resistance to infection are not completely understood, risk factors (such as HIV infection and history of previous TB) and markers associated with progression from Mtb infection to TB disease including predictive proteomic and gene-transcriptional signatures have been described [5-7]. Epidemiological risk factors such as proximity to index case, duration of employment, smoking and body mass index have not been directly associated with resistance, suggesting a potential role for cellular and/or genetic factors[1, 2, 8].

Resistance to Mtb infection has been shown to be relatively stable overtime in the HHC in Uganda ranging from 12% after 2 years to about 8% after about 8 years of follow-up [9]. Several factors may be associated with the durability of resistance to infection including frequency of exposure and the bacterial load of the index cases [10]. IGRA and TST are the only tests currently being programmatically used to diagnose Mtb infection or absence thereof. The limitations to these tests include the lack of specificity for Mtb for TST, and the reliance on a single cytokine (IFN-γ) for IGRA, as well as the use of only ESAT6 and CFP10 as antigens in the IGRA.

In our previous report from the same cohort, we tested a limited number of cytokines in QuantiFERON supernatants from the Nil tube or TB1 tube containing ESAT6/CFP10 peptides that preferentially induce CD4 responses. We showed that despite absence of IFN-γ in some of the individuals classified as not infected with Mtb (Resisters), there were detectable levels of other cytokines associated with T cell and innate cell responses [2]. Other studies have reported cytokine and antibody responses to the above antigens in the absence of IFN-γ responses, suggesting that some individuals who are IGRA negative may have Mtb infection but with alternative immune responses [9, 11, 12].

Here we used an expanded panel of immune markers to identify cytokine signatures that are associated with resistance to Mtb infection in response to live Mtb stimulation of PBMC. Characterising the resister phenotype and defining among them those who have true resistance to Mtb infection is critical for several reasons. It has the potential to enable a better understanding of the mechanisms which contribute to protection. Accurately defining and studying highly exposed resisters can reveal the epidemiological, genetic, and immunological factors that contribute to resistance, thus helping inform future strategies for prevention of TB infection including design of more effective vaccines through advancing knowledge of correlates of protection.

## Methods

### Study population

We conducted a cross-sectional study between January 2020 and December 2022 in which we enrolled HCW who had worked in high TB exposure hospitals or clinics for a minimum of 5 years, as previously described [2]. HCW were recruited from different TB hospitals and clinics or medical wards of public hospitals in the Western Cape province of South Africa. These sites included Brooklyn Chest Hospital, DP Marais Hospital, Site B and Site C clinics in Khayelitsha, Michael Mapongwana Community Health Centre, Mitchell’s Plain Hospital, Khayelitsha Hospital, Groote Schuur Hospital and Brewelskloof TB hospital. The study was approved by the University of Cape Town’s Faculty of Health Sciences Human Research Ethics Committee (HREC:031/2019). Adminisitrative approvals were also obtained from the Western Cape Department of Health and City of Cape Town.

After HCWs provided written consent, a screening questionnaire was administered to determine potential eligibility for enrolment into the study, and to collect epidemiological data from study participants including demographics and risk factors for TB infection. To qualify for enrolment into the study, participants had to be HIV seronegative, have no symptoms of active TB, no chronic obstructive pulmonary disease (COPD) or other respiratory illnesses, be older than 18 years and have worked in TB clinical settings for 5 years or more. Participants without any prior TB episodes were further screened for Mtb infection/exposure using IGRA (with the QuantiFERON TB Gold Plus kit) and TST. The IGRA was conducted according to the manufacturer’s instructions and IFN-γ concentrations (for both TB1 and TB2 tubes) < 0.35 IU/ml were considered negative and values ≥ 0.35 IU/ml were considered positive. TST was administered after blood was collected for IGRA to minimize any possible effect of TST on IGRA results, and indurations were read after 48-72 hours. Indurations <10 mm were considered negative, and indurations ≥10 mm were considered positive. Resisters were defined as IGRA <0.35 IU/ml and TST induration <10 mm. LTBI was defined as IGRA β 0.35 IU/ml and TST induration β 10 mm

### Extreme phenotypes

We further classified a subset of participants identified as Resisters or LTBI into groups of extreme phenotypes. The definition used for extreme Resisters was: IGRA <0.2 IU/ml and TST =0 mm; and extreme LTBI: IGRA β 1 IU/ml and TST induration ≥ 15 mm. Immunology assays described in this manuscript were conducted on participants with the extreme phenotypes.

### Sample collection and PBMC isolation

After classification into the different groups according to IGRA and TST status, participants were re-contacted and up to 50mL of whole blood was collected into sodium heparin tubes for plasma and PBMC isolation. A total of 2ml of blood was used for plasma separation by centrifugation at 1500 x g for 15 minutes. Plasma was collected and stored at −80°C and later used for evaluation of soluble markers as described below. PBMC were isolated from up to 48ml of whole blood using density gradient centrifugation with the Ficoll-Histopaque method. PBMC were cryopreserved in 10% fetal calf serum (FCS) and a final concentration of 10% dimethyl sulfoxide (DMSO) and stored overnight at −80°C, before being transferred to liquid nitrogen for long-term storage until required for analyses.

### PBMC stimulation

PBMC were retrieved from liquid nitrogen storage and thawed in a 37°C water bath. The cells were resuspended in 9 ml of pre-warmed R10 media and centrifuged at 400 xg for 10 minutes at room temperature (RT). Supernatants were discarded and pellets resuspended in 10 ml of R10 media, followed by overnight resting at 37°C in a 5% CO_2_ incubator. After resting, PBMC were centrifuged at 500xg for 5 minutes, and cell concentrations and viability were determined using trypan blue staining on a TC20 automated cell counter (Bio-Rad). PBMC were finally resuspended to a final concentration of 10 million cells/ml in preparation for stimulation assays.

### Antigens and stimulants

The following were used for PBMC stimulation: live H37Rv (Mtb) and live BCG (both used at MOI of 1, and cultured as described in the supplementary section), lipopolysaccharide (LPS, 100ng/mL; Invivogen), phytohaemagglutinin (PHA, 4 ug/mL; Remel).

### Flow cytometry

#### CD4 T cell panel

For stimulation assays, PBMCs were stimulated in BSL-3 laboratory with live Mtb, or live BCG. Phytohemagglutinin (PHA) and R10 media only were used as a positive and negative controls, respectively. For each stimulation condition, 100 µl of PBMCs (1 million cells) were transferred to 96 well plates, stimulated with 100 µl of either Mtb, BCG, PHA, or media only and incubated for 18 hours at 37°C. Following incubation, 100 µl of supernatant was collected without disturbing the cells, transferred to Sarstedt tubes, and stored at −80°C for use in Luminex assays. Stimulated PBMCs were then re-stimulated with the respective antigen cocktails in the presence of golgi-blockers (Monensin and Brefeldin A (BD Biosciences)) and incubated for an additional 6 hours. Plates were finally covered in aluminium foil and stored at 4°C in preparation for staining for flow cytometry. The following antibody-fluorochome combinations were used (Supplementary Table 1): CD3 FITC, CD4 BV785, CD8 APC-Cy7, Granzyme B BV421, IFN-γ Alexa Flour 700, TNF-α PE-Cy7 and the Live/Dead Dye (BV510).

#### Monocyte panel

Live Mtb, LPS, or R10 (media control) were used for stimulation in BSL-3 laboratory. For each stimulation condition, 100 µl of PBMCs (1 million cells) were transferred to 96 well plates, stimulated with 100 µl of antigens and incubated for 2 hours at 37°C in a 5% CO_2_ incubator. After the two hours, golgi-blockers (Monensin and Brefeldin A (BD Biosciences)) were added and incubated for an additional 6 hours. The following antibody-fluorochome combinations were used (Supplementary Table 1): CD3 ECD, CD14-PE-Cy7, HLA-DR BV605, TNF-α APC, IL-1β PE, and IL-6 FITC and near infra-red (NIR) viability marker (APC-Cy7).

Before staining, plates were first centrifuged at 500xg for 5 minutes to pellet cells, supernatants were discarded, and cell pellets washed with PBS and centrifuged again. First, cells were stained by adding 50 µl of the viability dye (resuspended in 50 µl DMSO and diluted with PBS) and incubated for 10 minutes at RT. Following incubation, samples were washed, centrifuged again and supernatants discarded. After viability staining, 50 µl of extracellular antibody cocktail containing phenotypic markers was added and samples incubated for 30 minutes at RT. This was followed by washing with FACS buffer and centrifugation to pellet the cells. Cells were permeabilized by adding 100 µl of Cytofix/Cytoperm (BD Biosciences) for 10 minutes at RT, washing with 100 µl of 1X PermWash Buffer (BD Biosciences). Intracellular cytokine staining was then conducted by adding 50 µl of staining antibody cocktail and incubating samples for 60 minutes at RT. Finally, cells were washed twice with PermWash buffer, and resuspended with 1% paraformaldehyde (PFA). Following staining, 500,000 events of each sample were acquired on a BD Fortessa cytometer using FACSDiva software. Flow cytometry data were analyzed with FlowJo version 9.9.6 (FlowJo LLC). For T cell and monocyte functions, background subtraction was carried out by subtracting the proportions in unstimulated cells from stimulated cells.

### Luminex assay

The following samples were used for this assay: plasma from whole blood, QuantiFERON TB Gold Plus supernatants (Nil and TB1 stimulated blood) and supernatants from PBMCs (Nil and stimulated with BCG and Mtb) collected as described above.

The ProcartaPlex Human Monitoring Panel 65-plex panel (Invitrogen™) (Supplementary material) was used and experiments were conducted according to the manufacturer’s instructions. Data was acquired on Bio-Rad Bio-Plex 200 Instrument with the acquisition of a minimum of 50 beads per analyte and analyzed using Bio-Plex Manager Software.

### Data and statistical analyses

All statistical analyses for Luminex were conducted on R software (R version: R 4.3.1) and its relevant associated packages [13-19]. The Wilcoxon test or Chi-squared test (χ2) or Fisher’s exact test were used for comparisons of categorical variables across study groups. Flow cytometry statistical analyses were performed using GraphPad Prism version 10. p-values <0.05 (with or without correction for multiple comparisons) were considered to be statistically significant.

### Luminex

All cytokines were included in the analyses. For markers with undetectable values, these were replaced with half the value for the lower limit of detection or the lowest extrapolated value for that marker. For markers with very high but undetectable values, these were replaced with twice the value for the upper limit of detection of the highest extrapolated value for that marker. For negative values after background subtraction (Mtb-Nil or BCG-Nil or TB1-Nil), the fold change was calculated (with group 1 as numerator and group 2 as denominator, depending on the analyses; either Resister vs LTBI or Resister_c2 vs Resister_c1) using the log2 of the absolute values of the fold change and then multiplied the result by −1 to maintain its direction. Statistical significance was assessed using Wilcoxon tests with False Discovery Rate (FDR, less than 0.05) for p-value adjustment.

### Dimensionality reduction from Mtb-stimulated PBMC supernatant samples

We applied a feature selection analysis with a random forest algorithm on the cytokine data from Mtb-stimulated PBMC (Mtb-Nil) as input separately, using caret and randomForest packages with 20-fold cross-validation and 5 repeats to assess model accuracy [20]. This analytical approach was applied to identify the most informative variables on the cytokine set. Then, the most predictive variables determined by the Mean Decreasing Gini and Mean Decreasing Accuracy indexes, were used in a generalized linear model to estimate the classification of LTBI and Resisters samples and the accuracy depicted using Receiver Operator Curve (ROC), in the plasma, PBMC and QuantiFERON samples, and the Area Under the Curve (AUC) with the pROC package.

### Identification of resister clusters

Resister sample results were used from Nil and Mtb-stimulated conditions and a delta condition was created by calculating Mtb-Nil for all cytokines. This delta table was used in the hierarchical clustering algorithm to define the resister subclusters, which were used as the main grouping variable for this analysis. Hierarchical clustering was performed using the Manhattan distance and Ward.D2 agglomeration method. After identifying two subclusters (henceforth labelled as Resisters_c2 vs Resisters_c1), we tested for differential abundances of cytokines between the clusters by applying Wilcoxon test, FDR and calculating log2 fold change for all the cytokines. Afterwards, the feature selection with random forest algorithm was applied to the differential cytokines to identify the most informative cytokines that would discriminate between the subclusters, using caret and randomForest packages, setting mtry = 1 and ntree = 400 as parameters, with the remaining parameters being maintained as default. The model identified the top 5 cytokines based on Mean Decreasing Gini and Mean Decreasing Accuracy indexes. Then, the 5-cytokines set were used in a generalized linear model to estimate the classification of the Resister subcluster samples. The 5-cytokines set’s overall accuracy was depicted using ROC analyses and its AUC values, in the plasma, PBMC and QuantiFERON samples after stimulation.

## Results

### Demographic characteristics

We enrolled a total of 780 participants who were HIV-uninfected and had worked for a minimum of 5 years (cumulatively) in a high TB exposure environment (Figure 1, Table 1). Of the 780 participants, 67 (8.5%) had TB treated in the past. A total of 659 had at least one test performed, and 100 participants either had IGRA test alone without the TST performed/measured, or an indeterminate IGRA test. A total of 559 had both IGRA and TST performed. Of the 559, 129 (23%) were double TST negative and IGRA negative fullfilling the criteria for Resisters, and 145 (26%) were TST positive and IGRA positive, fulfilling the criteria for LTBI for this study. The rest (285, 51%) had discordant results (positive for one test and negative for the other). From the 129 resisters and 145 LTBI, we further selected extreme Resisters (n=26) and extreme LTBI (n=24) (Table 1). We performed Luminex (Resisters, n=26; LTBI, n=18), and flow cytometry (LTBI n=14 and n=18 Resisters) on these participants with the extreme phenotypes.

**Table 1:**
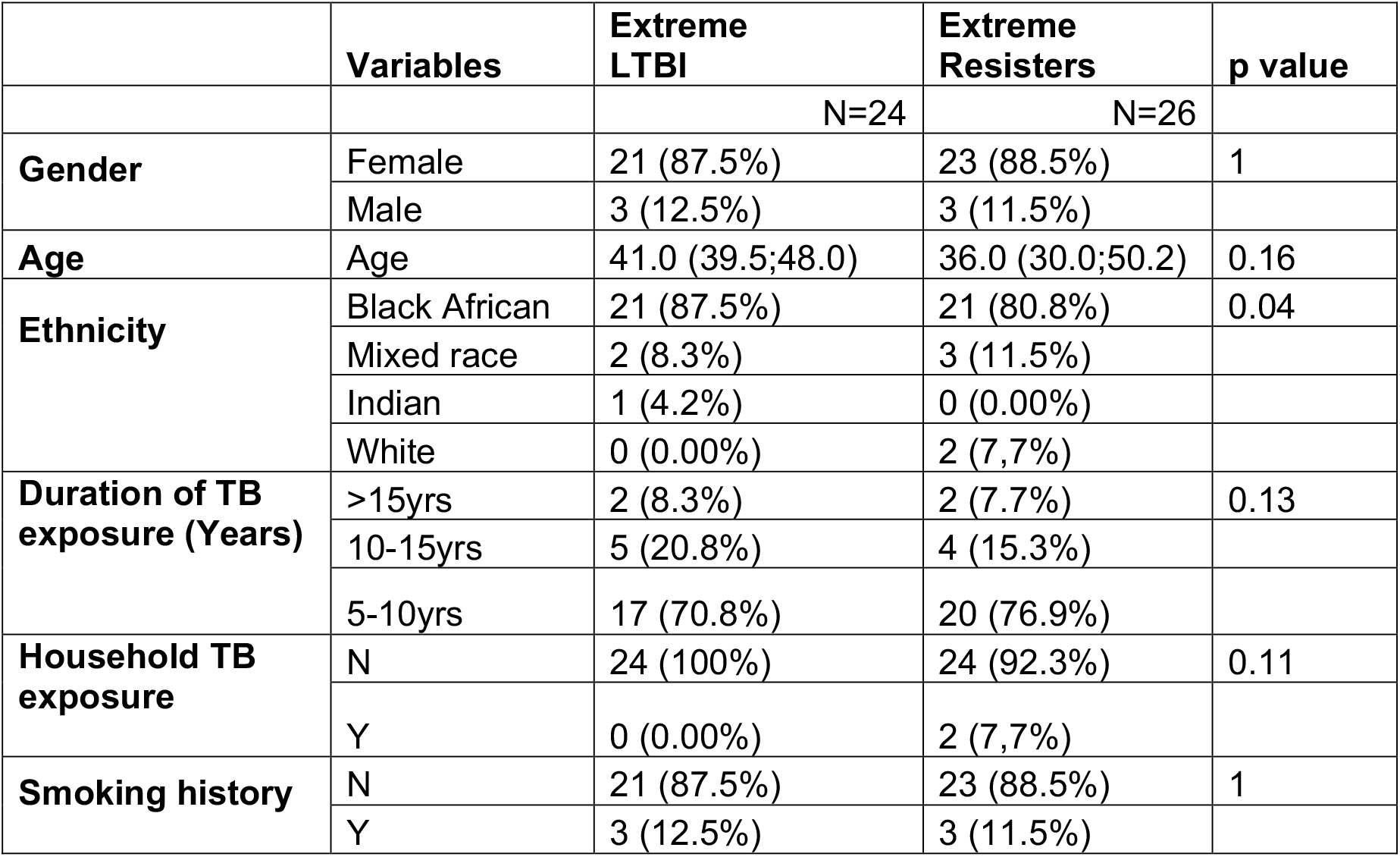
Demographic characteristics of all participants in the extreme Resisters and LTBI groups.

**Figure 1:**
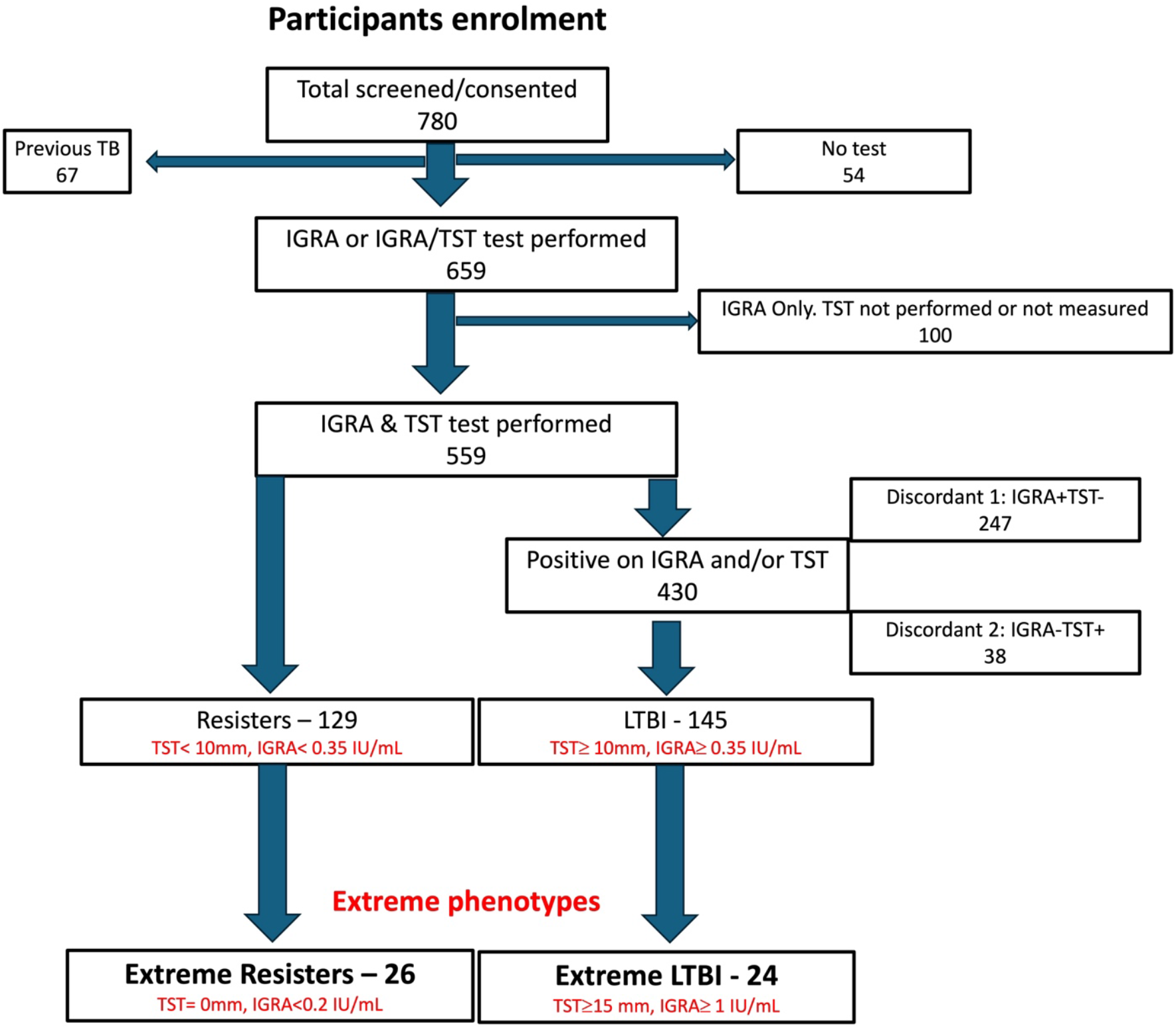
Schematic representation of the participant screening, enrolment and classification.

We compared the different variables collected between the extreme Resisters and extreme LTBI. As shown in Table 1, majority of the participants were female, of Black African ethnicity, worked between 5 and 10 years in a high TB exposure environment, had no household exposure over the past 20 years, and with no history of smoking. There were generally no statistically significant differences between the groups except for ethnicity, most likely driven by small numbers in the non-Black African group.

### Protein signatures differentiating Resisters from LTBI

From PBMC samples stimulated with Mtb, we used the variables selected by the random forest algorithm. The model presented an accuracy (Acc) of 1, 95% C.I. [0.974, 1], no information rate (NIR) of 0.5714, and the comparison of Acc > NIR was significant (p-value < 2.2×10-16). Based on the Mean decreasing Accuracy and Mean decreasing Gini indexes, the cytokines MCP-1 (CCL2), IL-17A, MDC and IL-8 (CXCL8) were the most informative for discriminating between LTBI and Resisters (Figure 2A). We tested the model variables in other samples to verify if the identified cytokines hold the same information in discriminating Resisters from LTBI. Thus, the value of MCP-1, IL-17A, MDC and IL-8 were retrieved from Plasma, PBMC (Mtb and BCG), and QuantiFERON (TB1). The markers presented similar performances in Plasma with AUC of 0.60 (CI: 0.45-0.75) (data not shown) and after stimulation, with AUC of 0.67 (CI: 0.54-0.85) on PBMC Mtb stimulation, 0.73 (CI:0.58-0.88) on PBMC BCG stimulation and 1 (CI: 1-1) on QuantiFERON TB1 stimulation (Figure 2B).

**Figure 2:**
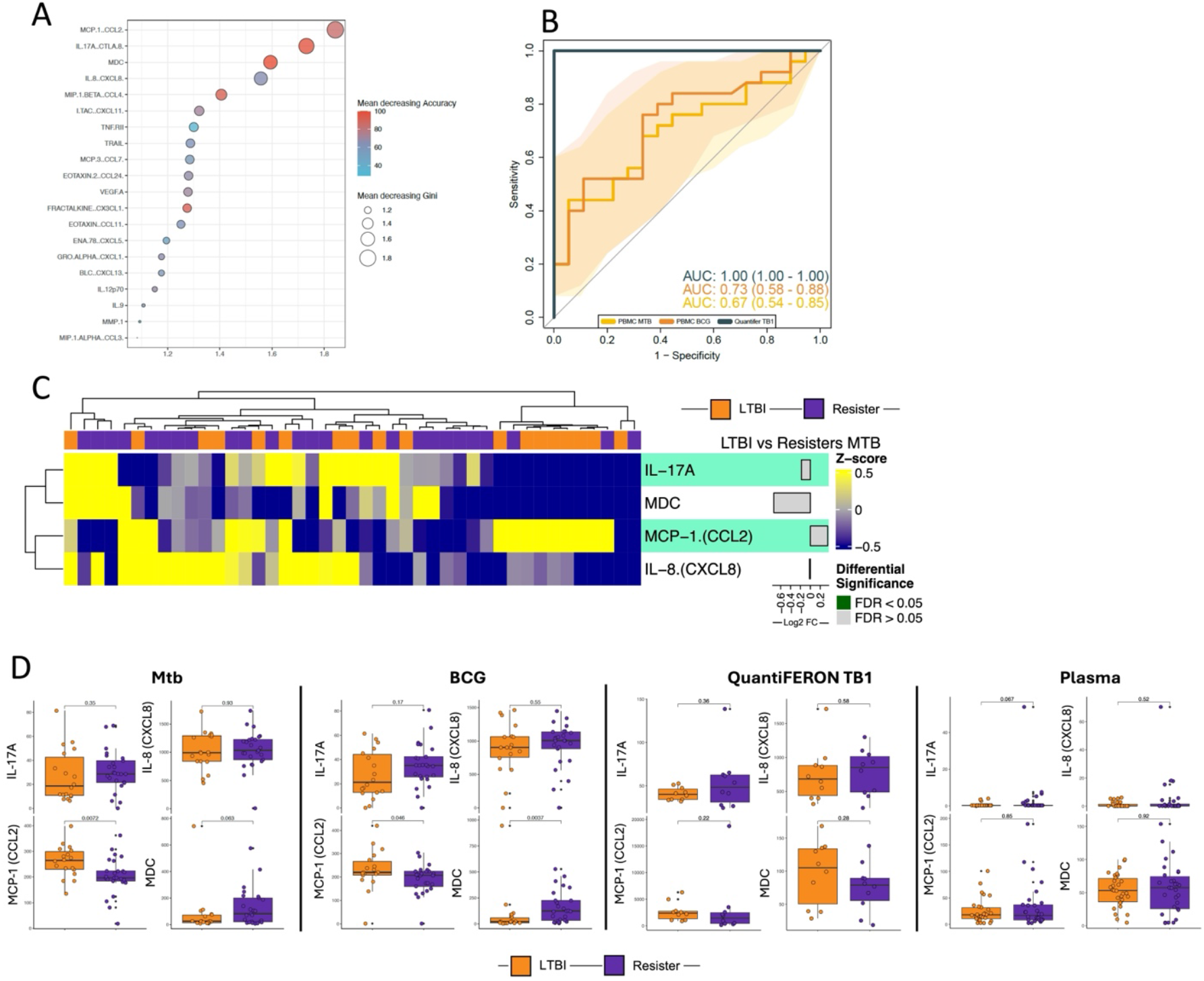
Identification of protein signature discriminating between Resisters and LTBI participants. A. Feature selection analysis identified markers discriminating between LTBI and Resister condition: Dot plot from the biomarker importance in the random forest model. Colors and dot size shows the Mean Decreasing Gene and Accuracy indexes from each variable. B. Receiver Operator Characteristics (ROC) analysis using the variables identified from dimensionality reduction concentrations to estimate the accuracy of the combined markers in discrimination of Resisters and LTBI with different stimuli. C. Heatmap of cytokines in protein signature from Mtb-stimulated PBMC. D. Boxplots of cytokines in protein signature in PBMC stimulated with Mtb and BCG, QuantiFERON stimulated with TB1 antigens, and plasma. Wilcoxon non-parametric test was applied to compare the LTBI and Resisters in each condition, without correction for multiple comparisons.

Next, we compared the abundance of the 4 cytokines between the Resisters and LTBI. Figure 2C shows the heatmap of the 4 cytokines in PBMC after Mtb stimulation. The concentrations of IL-17A were similar between the two groups across all stimulation conditions, but with a trend towards higher concentrations in Resisters in plasma ((p=0.067); Figure 2D). The concentrations of IL-8 were similar between the groups across all stimulation conditions and in plasma (Figure 2D). The concentrations of MCP-1 were similar between the groups in plasma and QuantiFERON samples, but significantly higher in LTBI after stimulation with both Mtb and BCG (p=0.0072 and p=0.046, respectively: Figure 2D). The concentrations of MDC were similar between the two groups for plasma and QuantiFERON samples, but significantly higher in Resisters after BCG stimulation (p=0.0037) and showed a trend towards higher concentration in Resisters after Mtb stimulation (p=0.063; Figure 2D).

### Mtb stimulation reveals clusters among the Resisters

We initially postulated that if there were to be any immunological differences between the Resisters and the LTBI participants, they were most likely to be seen with the extreme phenotype participants that were selected for these analyses. However, the differences we observed were not strong enough to withstand correction for multiple comparisons after FDR adjustment (data not shown). We therefore postulated that there may be heterogeneity in responses in the Resister group that could be masking potential differences between Resisters and LTBI. With this in mind, we focused our analyses on the Resister participants to characterise their immune response profiles.

Hierarchical clustering algorithm was applied to the data from the Resister PBMC samples stimulated with Mtb to identify clusters. Two clusters were identified within the Resister group: Resisters_c1 (n=10) and Resisters_c2 (n=14) (Figure 3A). After identifying the clusters, we compared the abundance of cytokines between Resisters_c2 vs Resisters_c1, which identified a total of 52 cytokines with differential abundance (37 upregulated and 15 downregulated in Resister_c2 compared to Resister_c1 (Figure 3A). Demographic characteristics were similar between the two clusters though Resister_c1 participants were all female (Table 2).

**Table 2:** Characteristics of participants in the two Resister clusters.

|  |  | Resister_c1 (n = 10) | Resister_c2 (n= 14) | p value |
| --- | --- | --- | --- | --- |
| Gender | Female | 10 (100%) | 11 (78.6%) | 0.239 |
|  | Male | 0 (0.00%) | 3 (21.4%) |  |
| Age (years) | Age | 37.0 (32.2 - 53.0) | 35.0 (26.2 - 47.2) | 0.305 |
| Ethnicity | Black African | 7 (70.0%) | 12 (85.7%) | 0.769 |
|  | Mixed race | 2 (20.0%) | 1 (7.14%) |  |
|  | Indian | 0 (0%) | 0 (0%) |  |
|  | White | 1 (10.0%) | 1 (7.14%) |  |
| Duration of TB exposure (years) | >15yrs | 1 (10.0%) | 1 (7.14%) | 1.000 |
|  | 10-15yrs | 1 (10.0%) | 2 (14.3%) |  |
|  | 5-10yrs | 8 (80.0%) | 11 (78.6%) |  |
| Household TB exposure | N | 9 (90.0%) | 13 (92.9%) | 1.000 |
|  | Y | 1 (10.0%) | 1 (7.14%) |  |
| Smoking history | N | 9 (90.0%) | 13 (92.9%) | 1.000 |
|  | Y | 1 (10.0%) | 1 (7.14%) |  |

**Figure 3:**
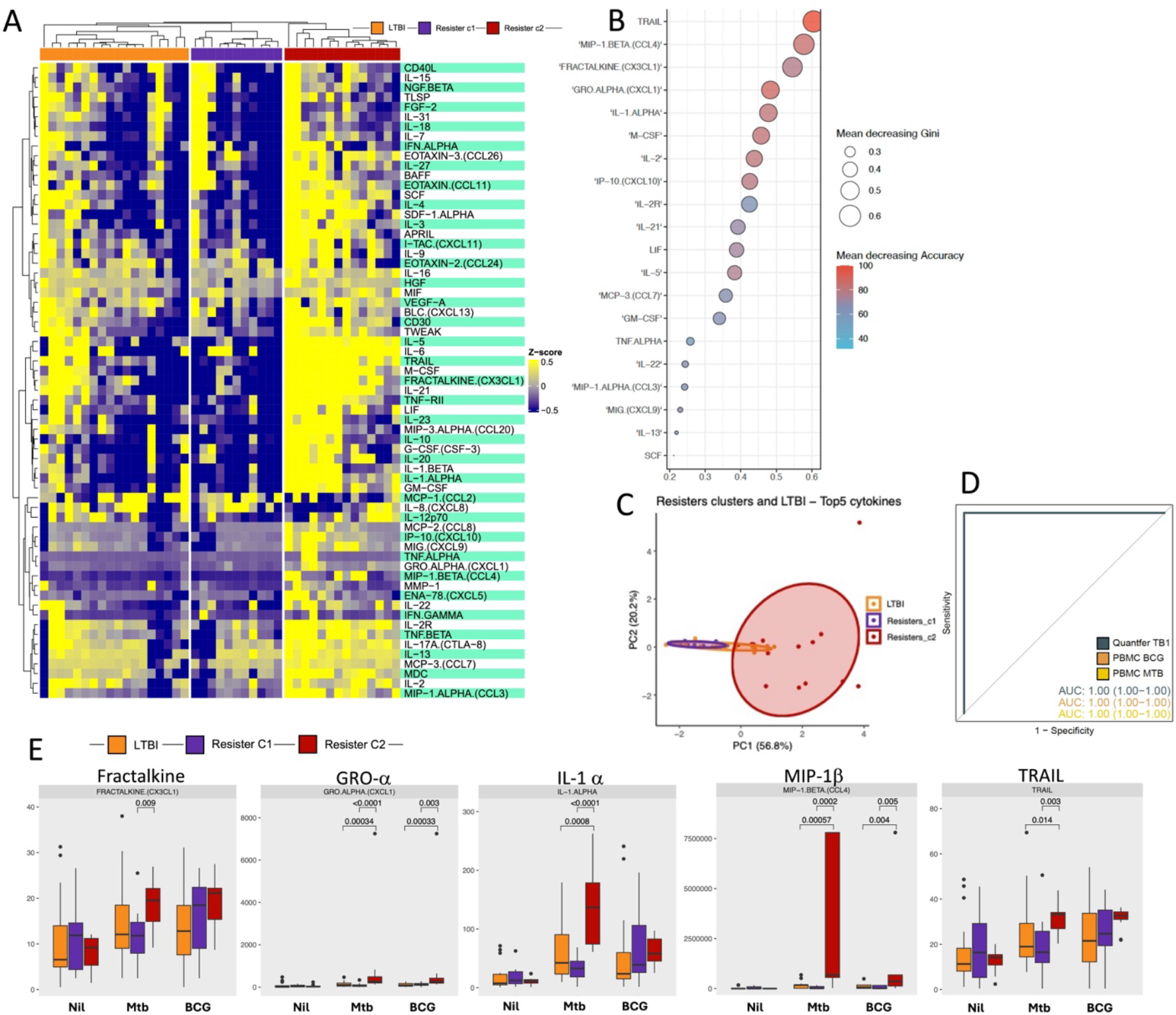
Identification of protein signature discriminating two distinct clusters in Resisters. A. Heatmap depicting overall cytokines abundance profile. Clusters were defined by hierarchical clustering using the Manhattan distance and Ward.D2 clustering methods. B. Feature selection analysis identified markers with higher importance and holding most of information regarding the Resister clusters. Dot plots from the biomarker importance in the random forest model. Colors and dot size shows the Mean Decreasing Gene and Accuracy indexes from each variable. C. PCA analyses discriminating the Resister clusters and LTBI. D. Receiver Operator Characteristics (ROC) analysis using the variables identified from dimensionality reduction concentrations to estimate the accuracy of the combined markers to discriminate Resister clusters with different stimuli. E. Boxplot displaying the most informative cytokines abundance values. Y axis represents the abundance values, while the X axis represents the different conditions. Wilcoxon non-parametric test, without correction for multiple comparisons, was applied to compare subclusters in each condition. The significance remained after correction for multiple comparisons, adjusting for FDR (data not shown).

We interrogated the cytokine data in order to identify protein signatures that may discriminate between the two Resister clusters from PBMCs stimulated with Mtb. Based on the Mean decreasing Accuracy and Mean decreasing Gini indexes, the cytokines TRAIL, MIP-1β (CCL4), Fractalkine (CX3CL1), GRO-α (CXCL1) and IL-1α combined together discriminated Resister_c1 and Resister_c2 (Figure 3B). The PCA analyses also revealed differences between Resister_c2 and Resister_c1, and similarity between Resister_c1 and LTBI (Figure 3C). We tested the model variables in the other samples to verify if the cytokines identified in the PBMC stimulated with Mtb performed similarly in discriminating the Resister clusters. In the plasma samples the AUC was 0.74 (CI: 0.53-0.94) (data not shown), and in the PBMC stimulated with BCG and MTB, and QuantiFERON stimulated with TB1 antigens, the AUCs were 1 (CI: 1-1) for all three sample types (Figure 3D).

Next, we compared the concentrations of the above five cytokines between the Resister_c2, Resister_c1, and the LTBI participants in PBMC samples stimulated with Mtb and BCG. All five cytokines were significantly higher in Resister_c2 compared with Resister_c1 after Mtb stimulation, but only GRO-α and MIP-1β were significantly different after BCG stimulation, still higher in Resister_c2 (Figure 3E). All five cytokines were also significantly higher in Resister_c2 compared with LTBI participants, but all were similar between Resister_c1 and LTBI for both Mtb and BCG stimulations (Figure 3E).

### Cellular responses in the Resister clusters

Based on the cytokine profiles between the Resister_c1 and Resister_c2 and based the similarity between Resister_c1 and LTBI, we hypothesized that Resister_c2 are those with true resistance to Mtb infection while Resister_c1 are those with alternative immune responses to Mtb infection. In addition, we hypothesized that the protein signature (with 5 cytokines) we identified that was higher in Resister_c2 (suggested to be true Resisters) is an immune signature of resistance that may be related to monocytes.

To check the first hypothesis, we analysed previously available data from the same participants in the Resister clusters (N=14 vs N=9) of CD4 T cells expressing IFN-γ, TNF-α and Granzyme B after stimulation with live Mtb and BCG (Figure 4A). We hypothesised that CD4 responses would be similar between LTBI and Resister_C1, and both higher than Resister_c2, suggesting that Resister_c1 but not Resister_c2 were Mtb infected. In response to both Mtb and BCG, we observed statistically similar frequencies of CD4 expressing IFN-γ, TNF-α and Granzyme B between the two Resister clusters and LTBI, albeit with a non-significant trend (p=0.076 and p=0.064) towards a decrease in frequencies of CD4 expressing TNF-α in Resister_c2 compared with Resister_c1 and LTBI, respectively, in response to BCG (Figure 4B).

**Figure 4:**
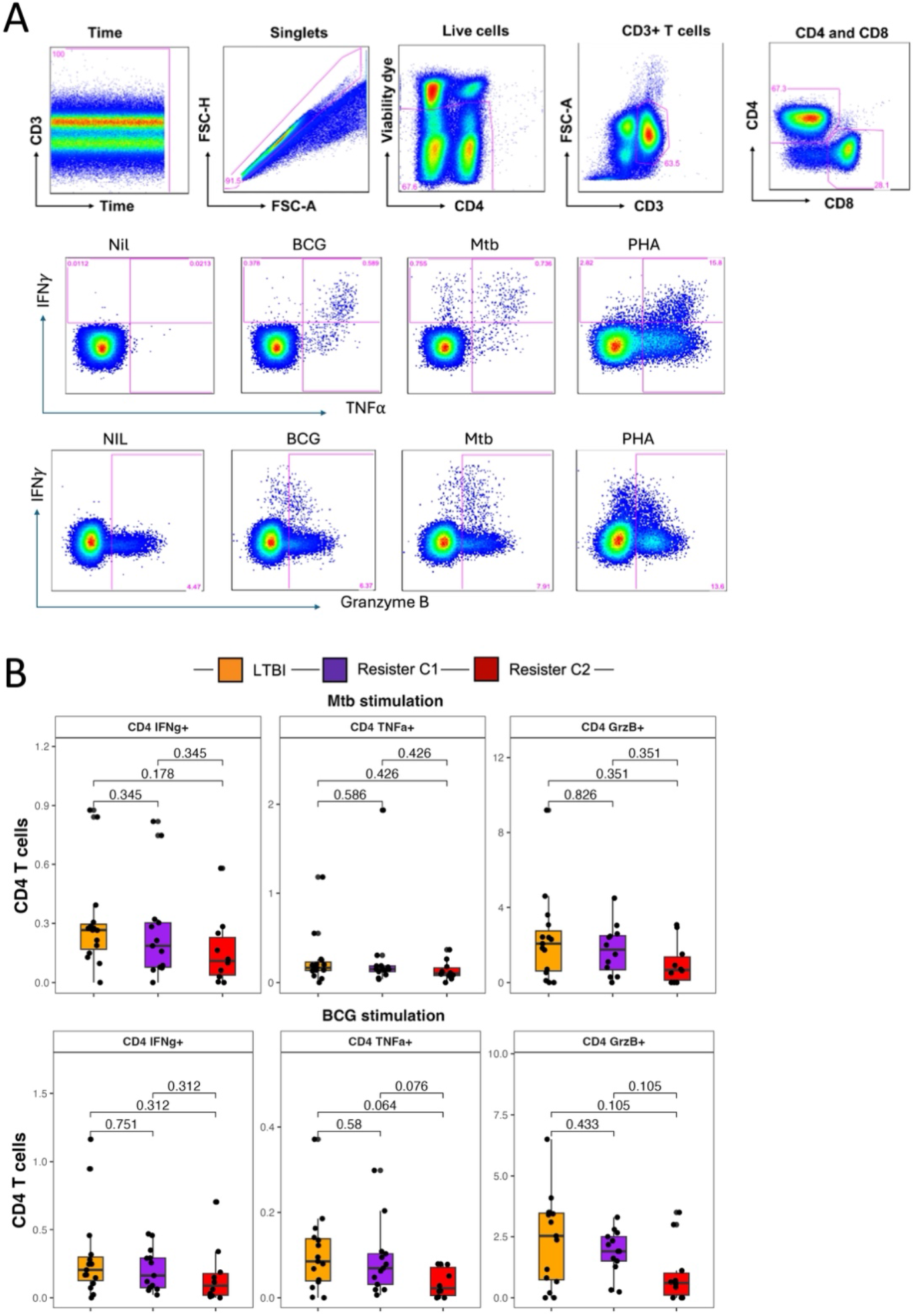
CD4 T cell responses in the Resister clusters and LTBI participants measured by flow cytometry. A. Simplified gating strategy for identification of CD4 T cells and the expression of intracellular cytokines and cytotoxic marker in response to Mtb, BCG and PHA. B. Summary data for the expression of the above markers after Mtb and BCG stimulation of PBMC in LTBI, Resister_c1 and Resister_c2. Wilcoxon non-parametric test was applied to compare the clusters in each condition, without correction for multiple corrections.

Next, we tested the second hypothesis that the protein signature was an immune signature of resistance related to myeloid cells, mainly monocytes. The hypothesis was that if this was the case, monocyte responses would be higher in Resister_c2 compared with the other two groups. To do this, we interrogated previously available data from the same participants in the Resister clusters from PBMCs that were stimulated with Mtb and LPS, and intracellular expression of IL-1β, IL-6 and TNF-α was measured. Of note, even though IL-1β, IL-6 and TNF-α were not part of the protein signature identified above, the levels of IL-6 and IL-1β were both higher (p<0.001) in Resister_c2 compared with Resister_c1, while TNF-α was similar across the groups from the Luminex experiments using PBMC stimulated with Mtb (Figure 3A). The frequencies of monocytes expressing all the cytokines separately or all three together were statistically similar between the groups for LPS and Mtb (Figure 5B).

**Figure 5:**
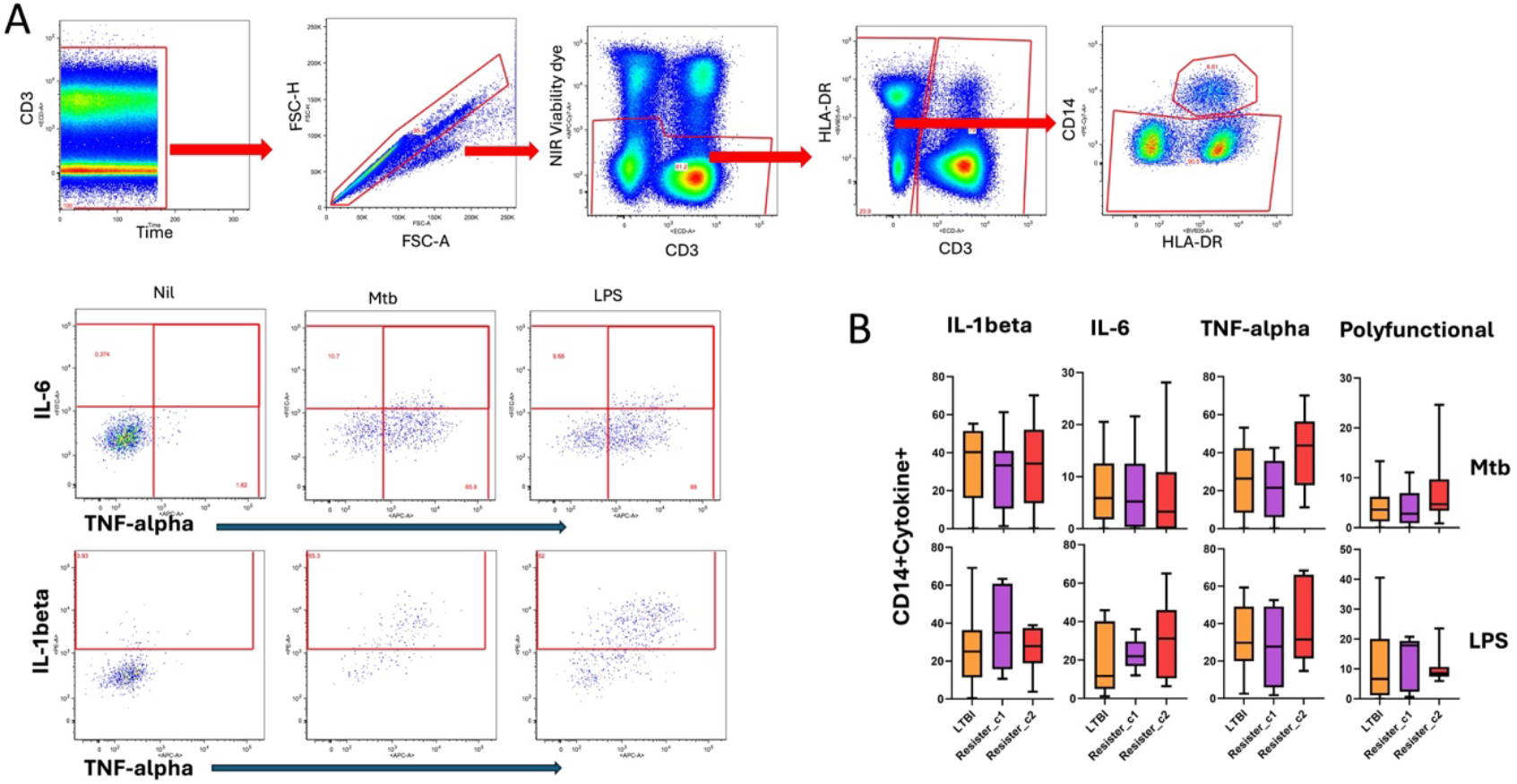
Monocyte responses in the Resister clusters and LTBI participants measured by flow cytometry. Simplified gating strategy for identification of monocytes and the intracellular expression of intracellular cytokines in response to Mtb and LPS. B. Summary data for the intracellular expression of cytokines after Mtb and LPS stimulation of PBMC in LTBI, Resister_c1 and Resister_c2. Wilcoxon non-parametric test was applied to compare the subclusters in each condition, without correction for multiple comparisons. All p values were >0.1 and thus not included in the figures.

## Discussion

We investigated factors associated with resistance to Mtb infection. We observed that 23% of the individuals screened and tested were classified as Resisters. In univariate analyses, there were no epidemiological factors that were different between the extreme Resisters and extreme LTBI participants used for immunological analyses. We identified a protein signature that discriminated between resisters and LTBI, consisting of IL-17A, IL-8, MCP-1 and MDC. Further analyses identified two resister clusters, with a combined protein signature of MIP-1β, TRAIL, IL-1α, Fractalkine and GRO-α, effectively discriminating the two clusters, with one cluster (Resister_c1) consistently having similar responses to LTBI, suggesting Mtb infection with alternative immune responses. The other cluster (Resister_c2) consistently had responses that were significantly different from Resister_c1 and LTBI, and Resister_c2 was thus hypothesized to be made of participants who have true resistance to Mtb infection.

The 23% of Resisters we observed was similar to the observations in a household contact study in Indonesia that tested and followed up participants for a period of 14 weeks to evaluate factors associated with early clearance of Mtb after household exposure [21], though this study used IGRA only to define infection with Mtb and the exposure period was much shorter than in our study. However, our percentage of Resisters was higher compared with the other studies in Uganda and South Africa on household contacts and goldminers that identified between 8-13% of participants being Resisters [3, 4, 9], even though some of these studies included both HIV infected and uninfected participants and used varying indurations of TST (0mm, 5mm, 10mm) to define TST positivity/negativity. The high prevalence of the resister phenotype could reflect a genuinely higher proportion of resisters in our high-exposure setting, possibly due to immune adaptation after sustained exposure. IGRA result reversions overtime during exposure could contribute to this observation, although we reasoned that sustained occupational exposure combined with dual testing using IGRA and TST would result in a relatively lower prevalence that what we observed. Longitudinal studies are needed to validate these findings and assess stability over time.

A protein signature consisting of IL-17A, IL-8, MCP-1 and MDC discriminated Resisters from LTBI with reasonable AUC values of more than 0.6 on ROC analyses. While there were no differences in IL-17A concentrations between Resisters and LTBI after stimulation, there was a trend towards a higher concentration in Resisters in plasma samples. A recent study including participants from household contact studies in Uganda reported that TH17-like CD4 T cells were associated with resistance where the proportions were higher in Resisters compared with LTBI after stimulation with Mtb antigens [22]. While we did not investigate cell associated IL-17A secretion, our data and others support the possible role for IL-17A/IL-17A-producing cells in protection from infection. The macrophage-derived chemokine (MDC) was observed to be higher in Resisters in response to bacterial stimulation. Though there have not yet been any reports directly implicating MDC in protection against Mtb infection, macrophages are widely known to play an important role in the initial immune response to Mtb in the lung.

Resistance to Mtb infection as currently defined by IGRA relies on IFN-γ production in response to ESAT6/CFP10. Based on published data [11, 12], we propose that resistance may exist in a spectrum with three possibilities: (1) some individuals are true Resisters (not infected, no antigen responses); (2) some individuals are infected but have IFN-γ-independent responses to Mtb antigens; (3) some individuals are infected but have IFN-γ responses to Mtb-specific antigens other than ESAT6/CFP10. Currently no published data exist that allows clear distinction between these three groups. Defining antigen specificity, or the type of immune responses in those who are actually infected among apparent Resisters will be critical for further research in this field and for developing strategies to appropriately manage these individuals such as prophylaxis to reduce the risk of progression to active disease for those individuals at higher risk, such as those with HIV infection and young children.

We identified two clusters within the Resisters and proposed that one of them may be true Resisters while the other is made of individuals with Mtb infection, mounting an alternative immune response to ESAT6/CFP10 or with responses to Mtb antigens other than ESAT6/CFP10. In this study we describe immunological markers that might reflect individuals with true resistance and alternative immune response to infection, with the latter having been suggested before in other studies including ours [2, 11, 22]. As current IGRA tests mainly rely on CD4 T cell memory responses to TB antigens as indication of an Mtb infection, we investigated CD4 responses to live Mtb but did not observe any significant differences between the Resister clusters. There was a trend towards lower frequencies of CD4 expressing TNF-α in Resister_c2 compared with Resister_c1 and LTBI, as hypothesized to indicate that Resister_c2 participants were true resisters. While concentrations of soluble markers showed significant differences, the lack of significant differences in CD4 responses between the clusters may be due to the fact that the sample size was not adequately powered to show cell-associated differences between the groups.

A protein signature consisting of MIP-1β, TRAIL, IL-1α, Fractalkine and GRO-α was identified to significantly discriminate between the two Resister clusters and could be a signature associated with true resistance, although further research is required to validate this signature and confirm its role as such. To our knowledge there exists no immunological marker that defines true resistance to Mtb infection after sustained exposure. A recent report from household contact study in Indonesia identified factors including BCG vaccination, innate immune cells numbers, innate cytokines, and antibodies that were associated with early clearance of Mtb after household exposure to a TB index case or after BCG vaccination of individuals with low Mtb exposure [21]. Innate immune cells such as granulocytes and monocytes were lower after 14 weeks of follow-up and after BCG vaccination, in those that remained persistently negative to IGRA test, while those that converted their IGRA status had no change between baseline and 14 weeks of follow-up. In addition, individuals with persistent negative results had higher cytokine production including IL-6 and IL-8 compared with converters and LTBI at baseline in response to *E. coli* stimulation. In this study, we observed a higher concentration of IL-6 and IL-1β in Resister_c2 compared with Resister_c1 in Mtb-stimulated PBMC, but we did not observe any significant differences between the resister clusters and LTBI in the frequencies of monocytes expressing either IL-6, IL-1β, TNF-α or a combination of all three in response to Mtb stimulation. The lack of significant differences may be due to small sample sizes not being adequate to show for cell-associated expression of markers. In addition, we did not investigate the cellular sources of the five cytokines that constituted the protein signature, and this may account for the lack of any observed significant differences in the CD4 and monocyte cytokine responses by flow cytometry.

The strengths of our study include the fact that we used both TST and IGRA to define Mtb infection as well as the use of samples with IGRA/TST values at the opposite end of the spectrum for the definition of extreme LTBI and extreme Resisters, thereby eliminating the possibility of misclassification of participants especially those with borderline results of the two tests. In addition, the medians and interquartile ranges for the cytokines in the protein signature defining the two Resister clusters do not overlap giving strength to the observations. Limitations include lack of a control group of participants without exposure and infection with Mtb. In this study, despite sustained exposure, the intensity of the exposure could not be well quantified. The other limitation is that we did not evaluate the responses in the resister clusters to other Mtb peptides or conduct longitudinal sampling to evaluate longevity of resistance.

In conclusion, we showed that cytokine responses to Mtb segregate into two distinct clusters among individuals classified as Resisters after prolonged Mtb exposure in health care facilities. We hypothesise that one of these clusters represents individuals who have true resistance to Mtb infection. The protein signature associated with this cluster needs to be further evaluated and could potentially define natural immunity to infection.

## Author contributions

MS conceived the study, obtained funding, wrote all protocols and SOPs, processed and stored samples, performed laboratory experiments and analyses, supervised all aspects of the project and wrote the first draft of the paper. Aba and NM (contributed equally to this work) processed and stored samples, performed the experiments and analyses, reviewed the manuscript. Abe processed and stored samples and reviewed the manuscript. CS provided clinical and scientific inputs throughout the study, assisted with data analyses and reviewed the manuscript. RG was involved in project management, and reviewed the manuscript. KAW, DaL, and DeL provided critical scientific, clinical and data analysis input and reviewed the manuscript. ATLQ, ERF and BBA did the bioinformatics analyses and reviewed the manuscript. GM was involved from project conception, provided critical scientific and clinical inputs, mentorship and reviewed the manuscript. All authors contributed to the article and approved the submitted version.

## Acknowledgements

We thank all the participants who participated in the study, Ms Rachel Dielle (Study nurse) and Ms Fuzeka Mdlulwa (Clinical Research Worker) who assisted with the consenting, recruitment of study participants and collection of biological samples.

