## Supplementary material for "Two distinct cytokine response clusters identified in healthcare workers with apparent resistance to infection with *Mycobacterium tuberculosis* despite sustained occupational exposure"

**Supplementary Table 1**: Antibodies and fluorochrome combinations used in the flow cytometry panels

| Antibody | Fluorochrome /Channel | Clone | Supplier | Volume in 50 µL |
|---|---|---|---|---|
| **CD4 panel** | | | | |
| Live/Dead Dye | BV510 | Aqua fluorescent reactive dye | Invitrogen | 0.5 |
| CD4 | BV785 | SK3 | BioLegend | 0.6 |
| HLA-DR | PE | L243 | BioLegend | 1 |
| CD3 | FITC | DREG-56 | BioLegend | 1 |
| CD8 | APC-Cy7 | SK1 | BioLegend | 2 |
| Granzyme B | BV421 | GB11 | BioLegend | 4 |
| TNF-$\alpha$ | PE-Cy7 | MAb11 | BioLegend | 1 |
| IFN-$\gamma$ | Alexa Flour700 | B27 | BioLegend | 0.5 |
| **Monocyte panel** | | | | |
| NIR | APC-Cy7 | Near Infrared Dye | Invitrogen | 1 |
| CD14 | PE-Cy7 | M5E2 | ELabscience | 2,5 |
| CD3 | ECD | A07748 | Beckman Coulter | 1,2 |
| HLA-DR | BV605 | L243 | BioLegend | 0.5 |
| TNF-$\alpha$ | APC | R101 | Sinobiological | 2.5 |
| IL-6 | FITC | MQ2-13A5 | ELabscience | 1.2 |
| IL-1$\beta$ | PE | AS10 | BD | 0.6 |

**Supplementary information**

**BCG culture**

Bacillus Calmette-Guerin (BCG) is the only licensed vaccine for TB and is a TB strain made up of live *Mycobacterium bovis*. BCG shares several proteins with Mtb but lacks genes from the region of interest (RD-1) which confer virulence to Mtb (23).

We cultured BCG stocks to exponential growth phase in 7H9 media supplemented with glycerol, tween-80, and Oleic Albumin Dextrose Catalase (OADC). Once cultures reached the exponential growth phase (OD value between 0.4 and 0.7), stocks were made by storing the cells in 7H9 media with 15% glycerol and transferring cells to -80°C for long-term storage. Frozen stocks were used to determine colony forming units (CFU) by plating different dilutions of thawed BCG in 7H10 agar plates and incubating plates at 37°C. Cells were counted after 3 weeks of incubation. BCG was used at MOI of 1.

**H37Rv culture**

Mtb H37Rv was cultured in the BSL-3 laboratory in 7H9 media supplemented with glycerol, tween 80, and OADC until culture reached exponential growth phase. Clumps were dispersed using glass beads and bacterial stocks were prepared by storing cells in

7H9 media supplemented with 15% glycerol. To determine CFUs, different dilutions of thawed bacterial stocks were cultured in 7H10 agar plates and incubated at 37°C for 3 weeks. Colonies were manually counted to determine CFUs. H37Rv was used at MOI of 1.

**List of cytokines used in Luminex and bead regions**

APRIL [52], BAFF [65], BLC (CXCL13) [15], CD30 [34], ENA-78 (CXCL5) [55], Eotaxin-2 (CCL24) [37], Eotaxin-3 (CCL26) [77], FGF-2 [75], Fractalkine (CX3CL1) [39], IL-16 [47], IL-2R (CD25) [22], IL-20 [26], I-TAC (CXCL11) [46], MCP-2 (CCL8) [18], MCP-3 (CCL7) [66], MDC (CCL22) [30], MIF [67], MIG (CXCL9) [54], TNF-RII [45], TRAIL (CD253) [29], TSLP [61], TWEAK [35], CD40L (CD154) [74], Eotaxin (CCL11) [33], Gro-alpha (CXCL1) [61], G-CSF (CSF-3) [42], GM-CSF [44], HGF [46], IFN alpha [48], IP-10 (CXCL10) [22], IFN gamma [43], IL-1 alpha [62], IL-1 beta [18], IL-2 [19], IL-3 [73], IL-4 [20], IL-5 [21], IL-6 [25], IL-7 [26], IL-8 (CXCL8) [27], IL-9 [52], IL-10 [28], IL-12p70 [34], IL-13 [35], IL-15 [65], IL-17A (CTLA-8) [36], IL-18 [66], IL-21 [72], IL-22 [76], IL-23 [63], IL-27 [14], IL-31 [37], LIF [15], M-CSF [67], MCP-1 (CCL2) [51], MIP-1 alpha (CCL3) [12], MIP-1 beta (CCL4) [47], MIP-3 alpha (CCL20) [56], MMP-1 [64], NGF beta [55], SDF-1 alpha (CXCL12) [13], SCF [39], TNF alpha [45], TNF beta [54], VEGF-A [78]
